# Classifying Biophysical Subpopulations of Insulin Secretory Granules using Quantitative Whole Cell Structure Analysis

**DOI:** 10.1101/2025.09.09.675239

**Authors:** Kevin Chang, Aneesh Deshmukh, Riva Verma, Valentina Loconte, Kate L. White

## Abstract

Pancreatic β-cells contain insulin secretory granules (ISGs), organelles where proinsulin is converted into insulin. As ISGs mature, they undergo extensive biophysical remodeling, producing a spectrum of subpopulations with heterogeneous molecular and spatial characteristics. However, systematic methods to define ISG subpopulations remain underdeveloped. To address this gap in knowledge, we employed soft x-ray tomography (SXT), which can quantitatively measure the biochemical density of ISGs within whole β-cells. Using unsupervised clustering, we classified subpopulations based on molecular density, size, and spatial positioning. Across different insulin secretory stimuli, we observed shifts towards mature and releasable subtypes, demonstrating that exogenous signals can dynamically remodel ISG subpopulation distributions. We extended this methodology to primary β-cells characterized using volume electron microscopy (vEM). Integrating subpopulations from SXT and vEM uncovered insights inaccessible by a single method in isolation. This strategy establishes a framework for defining therapeutic approaches aimed at enriching physiologically beneficial ISG subpopulations.

## INTRODUCTION

Pancreatic β-cells secrete insulin, the central hormone responsible for blood glucose homeostasis.^1^ Within β-cells, active insulin is converted from proinsulin inside organelles called insulin secretory granules (ISGs).^2^ During this process, known as ISG maturation, the ISG interior acidifies and insulin molecules crystallize into a dense core.^3,4^ The maturation of ISGs results in a heterogeneous population of organelles characterized by differences in pH, protein and lipid composition, age, motility, and release state.^4–11^ Notably, defects in ISG maturation and differences in ISG subpopulation composition are implicated in both type 1 and type 2 diabetes.^12,13^ To capture distinctions related to exocytosis, ISG subpopulations are commonly classified into the readily releasable pool (RRP) and reserve pool.^14^ While this definition is useful for understanding release competency, it does not encompass the full range of ISG heterogeneity such as insulin processing state or ISG remodeling due to changes in the chemical environment.^15^ Therefore, a more systematic and unbiased characterization of ISGs based on multiple parameters would improve our understanding of ISG subpopulations. Furthermore, understanding the effect of current therapeutics and key insulin secretory signaling pathways on the distribution of ISG subpopulations may offer new insights for developing novel therapeutic strategies that enrich specific physiologically beneficial subpopulations.

A comprehensive method to classify ISG subpopulations would be based on multiple biophysical parameters and sample ISGs from whole cells.^16,17^ Previous efforts have leveraged isolated ISGs for proteomic, lipidomic, and biophysical characterization to identify subpopulations.^7,10,18^ While these approaches are useful for identifying changes in ISG composition under different experimental conditions, isolating ISGs from cells removes the cellular context of subpopulation identity. To address these constraints, we sought to use a method that can quantify ISG characteristics in whole cells.

A pioneering technique to study the biochemical content of individual ISGs within whole β-cells is soft x-ray tomography (SXT).^19–21^ In SXT, cells undergo cryofixation and are imaged using a full rotation stage, enabling the collection of whole volume, near native state data.^22^ Additionally, cells are imaged at the “water window” (284 to 543 eV), where biochemically rich material has a much higher relative absorbance than the surrounding cytoplasmic background. The x-ray absorption can be expressed as the linear absorption coefficient (LAC) parameter, which quantifies the molecular density of organelles such as ISGs.^23^ Previous studies have demonstrated the utility of the LAC parameter for investigating ISG maturation at the scale of individual granules.^24^ This approach is also relatively high-throughput, enabling the collection of larger datasets with multiple stimulation conditions for analysis compared to other 3D whole cell approaches. Therefore, SXT can offer new insights into ISG subpopulations by revealing the chemical state of ISGs across diverse cellular states.

To meaningfully categorize ISG subpopulations and understand how specific cellular stimuli impact subpopulation profiles, we applied unsupervised clustering on the chemical and physical features of ISGs such as morphology, molecular density, and position in the cell. In the field of quantitative cell biology, unsupervised clustering has been typically used on transcriptomic data to classify subpopulation of cells, but recently has also been employed in diverse cellular imaging methods to find inherent groupings of proteins and organelles without explicit labels.^25–28^ We applied clustering analysis to a pre-existing SXT datasets of INS-1E rat insulinoma cells and mouse primary β-cells generated to study ISG maturation.^24^ Exposure to orthogonal insulin secretion stimuli led to significant biophysical remodeling of ISGs, making this dataset a valuable resource for studying ISG subpopulations. We find that the classified ISGs subpopulations reflect a variety of insulin maturation states at different cellular spatial zones. This clustering methodology also lets us understand how different insulin secretagogues shift ISG subpopulations into more processed, mature ISG states. To extend this approach, alternative clustering methodologies were used to explore additional facets of ISG heterogeneity.

Another approach to exhaustively sample ISGs from whole cells would be volumetric electron microscopy (EM).^29^ While lacking quantitative information about biochemical density, this family of methods has been shown to accurately measure ISG characteristics at high resolution such as subtle associations between ISGs and microtubules.^30^ To explore the generalizability of our approach and complement our SXT-based subpopulations, we simultaneously performed subpopulation analysis using a high-resolution volumetric EM dataset obtained from mouse primary β-cells.^30^ This work outlines a new methodology to classify ISG subpopulations from diverse whole cell datasets and understand how different stimulatory conditions can increase ISG subpopulations of potential therapeutic interest.

## RESULTS

### SXT provides multiparameter descriptions of ISGs

Using the full rotation SXT method, we were able to image the entire volume of a β-cell (Figure 1A). The LAC-based contrast mechanism of SXT allows us to identify a range of ISG maturities (Figure 1B).^31^ Tomogram segmentations resulted in label fields of key organelles such as the ISGs, mitochondria, nucleus, and plasma membrane (PM) across whole cells (Figure 1C).

**Figure 1.**
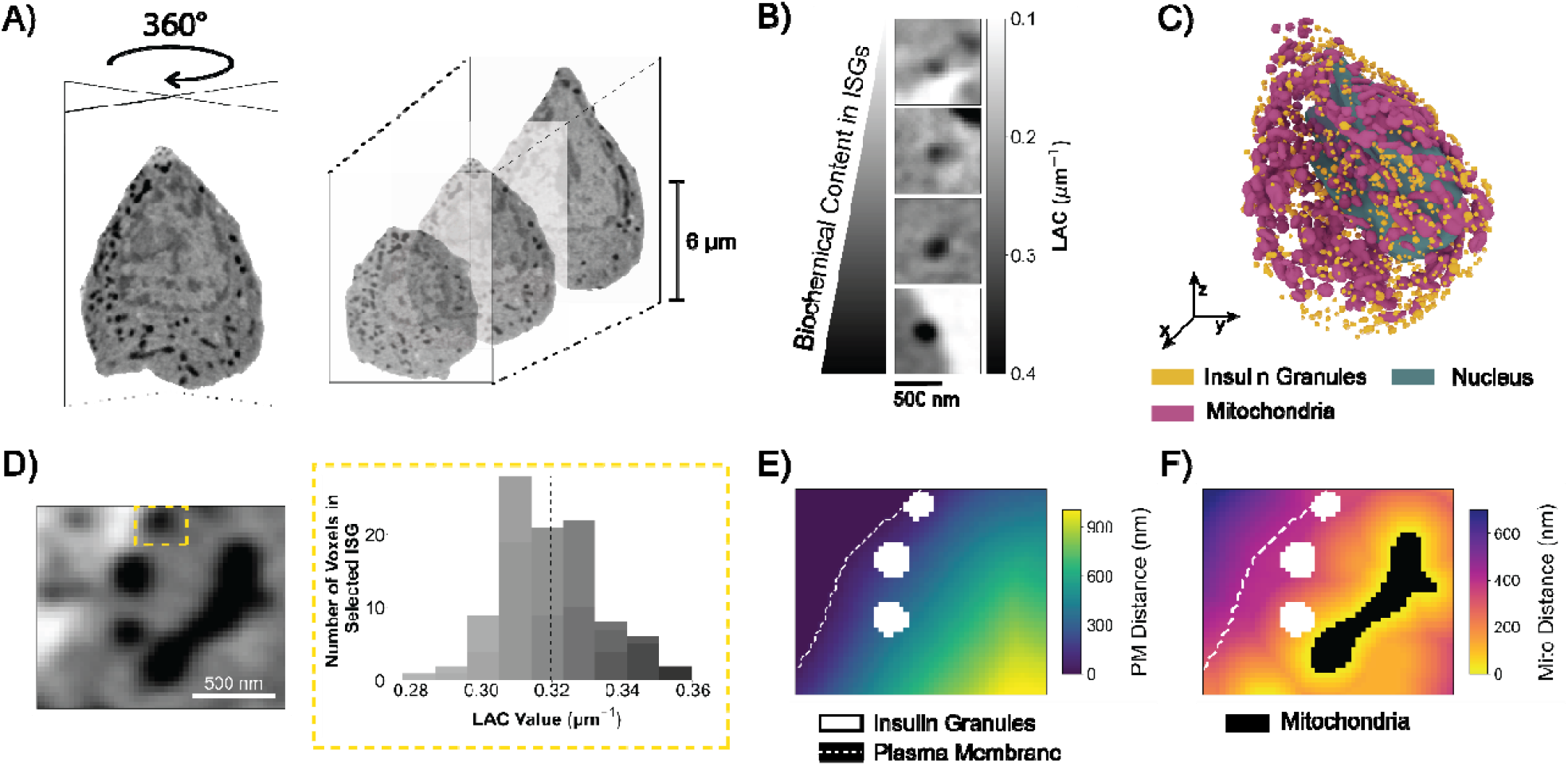
SXT imaging and calculation of ISG parameters. (A) SXT full-rotation projection images, which are combined to create a 3D image stack. (B) The LAC values in the SXT data correspond to biochemical content and are necessary for quantifying ISG maturities. (C) Blender rendering of a segmented INS-1E cell. ISG, nucleus, and mitochondria masks are shown. (D) Zoomed in image of an SXT orthoslice and LAC histogram of a selected ISG. Analysis at this resolution enables the creation of individual granule LAC statistics. (E) ISGs overlaid on the EDT of the plasma membrane. ISGs are displayed in white and PM displayed as a dashed white line. (F) Mitochondria mask inverse EDT with overlaid organelles. ISGs are again displayed in white, with PM displayed as a dashed white line. Mitochondria are colored black.

Individual ISGs were parameterized by their LAC values, size, and distances from other important organelles (Figure S1A). Each ISG has a distribution of LAC values (Figure 1D), which allows for the calculation of a LAC mean, standard deviation, skew and kurtosis (Figure S1B). To account for the ellipsoid shape of some granules, the geometric mean diameter was used to represent ISG diameters. We computed comprehensive distances between ISGs and cellular structures like the mitochondria and PM by using the Euclidean distance transformation (EDT) of the corresponding organelle masks (Figure 1E, F, S1C). Distance from the PM is a key parameter, since it provides insight into the mobilization of ISGs for secretion. These approaches collectively yield a set of parameters used to identify unique ISG subtypes: LAC mean, standard deviation, skew, kurtosis, diameter, and distances from the PM and mitochondria (Figure S2).

### Clustering framework to define ISG subpopulations

After establishing a list of parameters for each ISG, we set out to create a comprehensive unsupervised clustering methodology to classify ISG subpopulations. We reasoned that groupings of ISGs should remain consistent even if the underlying data is perturbed in a non-essential manner such as bootstrapping.^32^ To implement this reasoning, cluster stability analysis was used. This analysis validates that over many bootstrapped datasets, the cluster assignments for individual ISGs are relatively stable as measured by the Jaccard coefficient.^33^ Additionally, to create generalizable ISG subpopulations, we used multiple cells to average over individual cell variation and responses to experimental conditions. In total, ∼36,000 ISGs from 44 INS-1E cells in seven experimental conditions were analyzed. We aggregated ISGs from all the cells and performed clustering on this pooled dataset to create our final subpopulation assignments. In this approach, subpopulations are composed of ISGs with similar characteristics across multiple INS-1E cells.^34^

After applying stability analysis and k-medians clustering on the aggregated dataset, we found 5 ISG subpopulations (Figure S3A). ISGs were visualized and labeled by their assigned subpopulation within multidimensional uniform manifold approximation and projection (UMAP) space (Figure 2A). When ISGs are colored by their individual parameter values such as LAC mean, UMAP illustrations enable association between specific ISG features and subpopulation identity (Figure S3B). Subpopulations were labeled in order of increasing average LAC mean to reflect relative ISG maturity (Figure 2B). To further characterize these subpopulations, we determined that distance from the PM and LAC mean were more important to subpopulation identity than LAC skew and kurtosis (Figure S4). Additionally, one potential limitation of grouping ISGs across cells is the risk of misrepresenting individual cells. To validate the applicability of this approach, we found that these subpopulations broadly capture the characteristics of each INS-1E cell in our dataset (Figure S5).

**Figure 2.**
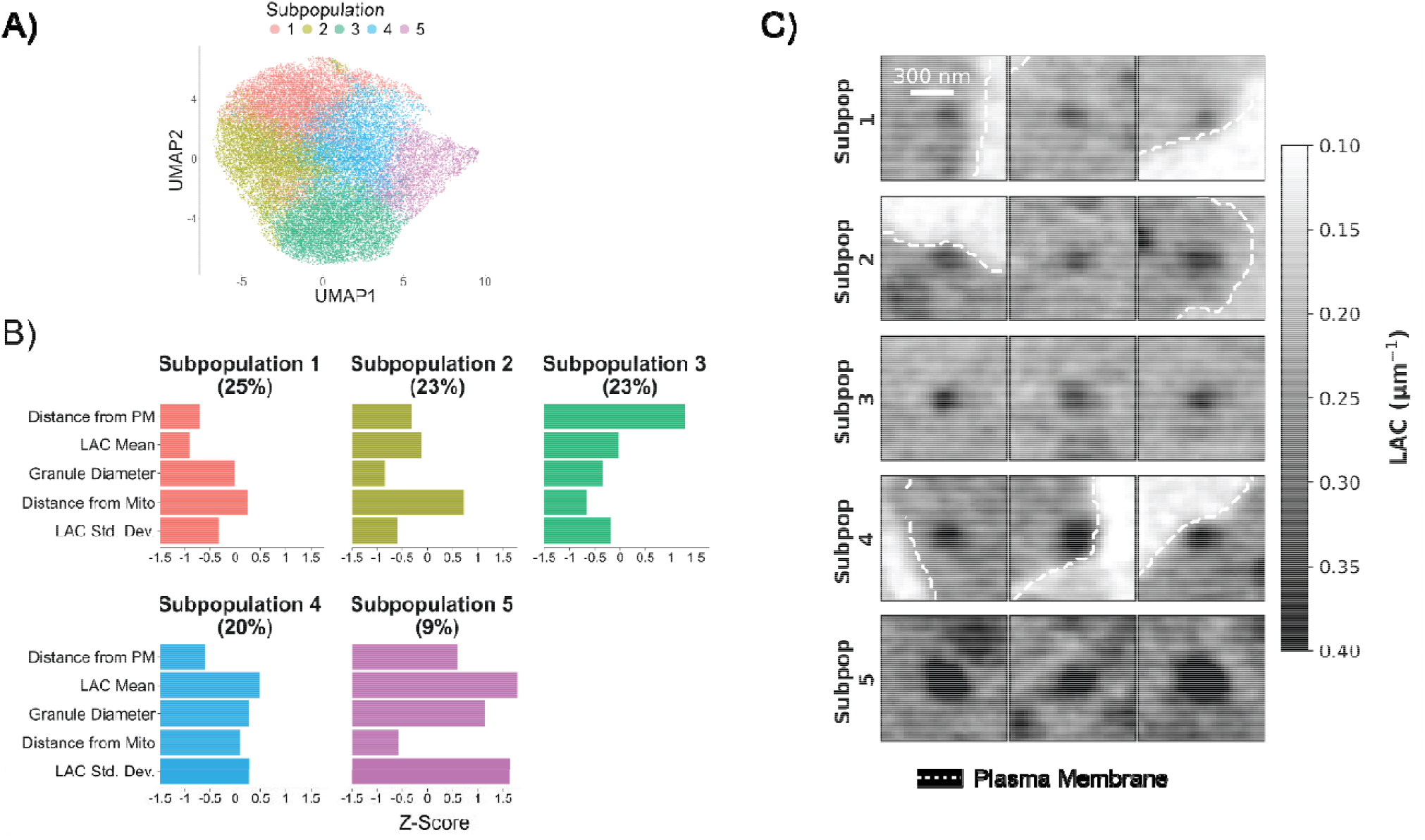
Visualizing ISG subpopulations in INS-1E cells. (A) UMAP diagram displays a dimensional reduction of individual ISGs from our dataset. ISGs are colored by their subpopulation identity assignment. (B) ISG Subpopulation profile plot. Summarized parameters are calculated using the median Z-score. Subtitles indicate the percentage of subpopulations out of the total number of ISGs. The subpopulations are numbered according to their LAC Mean, while parameters are listed from top to bottom in order of importance to subpopulation identity. (C) Gallery of ISGs from the five subpopulations within SXT projection images. PM displayed as a dashed white line.

We interpret Subpopulation 1 as an immature pool of ISGs primarily due to their reduced LAC mean (Figure 2B). Although ISGs with minimal biochemical density may go undetected using SXT, integrated structural models of granules at earlier maturation stages suggest they still possess appreciable LAC content.^31^ While Subpopulation 2 ISGs have smaller diameters than Subpopulation 1 ISGs, they exhibit an intermediate LAC mean and reduced LAC standard deviation phenotype. This suggests that these are condensed granules with a large core size relative to total granule size. Subpopulation 3 is far from the PM, so this subpopulation may represent the internal reserve pool of ISGs.^35^ Subpopulation 4 is closer to the plasma membrane and has relatively high LAC and above average diameter. This pattern was observed in a previous cryo-electron tomography (cryo-ET) study, where the authors found a greater proportion of larger, mature ISGs near the cell periphery.^36^ Therefore, these granules are likely mature ISGs with a PM localization comparable to the readily releasable pool. The last grouping, Subpopulation 5, is notable since it is far from the plasma membrane yet contains ISGs with very high LAC mean, larger diameter, and elevated LAC standard deviation (Figure 2B,C). Increasing the display range of maximum LAC illustrates the high LAC standard deviation of Subpopulation 5 ISGs, indicating that these granules occupy a unique state in the ISG maturation process (Figure S6). As a result, we consider this subpopulation to be a secondary, smaller group (9% of total ISGs) of mature and highly dense ISGs held in reserve.^3^ While it is possible that lysosomes could be mislabeled as large ISGs, prior correlative cryo-fluorescence analysis indicates that potentially mislabeled lysosomes would make up a negligible proportion of the overall dataset.^24^

### β-Cell stimulation conditions remodel ISG subpopulation distributions

To understand biologically meaningful changes in ISG subpopulations, we analyzed ISG subpopulations across six stimulation conditions associated with different cellular signaling pathways along with no stimulation as a control (Figure S7). Grouping by experimental condition reveals that different insulin secretagogues shift the relative distribution of ISG subpopulations (Figure 3A). This finding is visually apparent when ISGs in representative cells from each condition are colored by subpopulation identity (Figure 3B).

**Figure 3.**
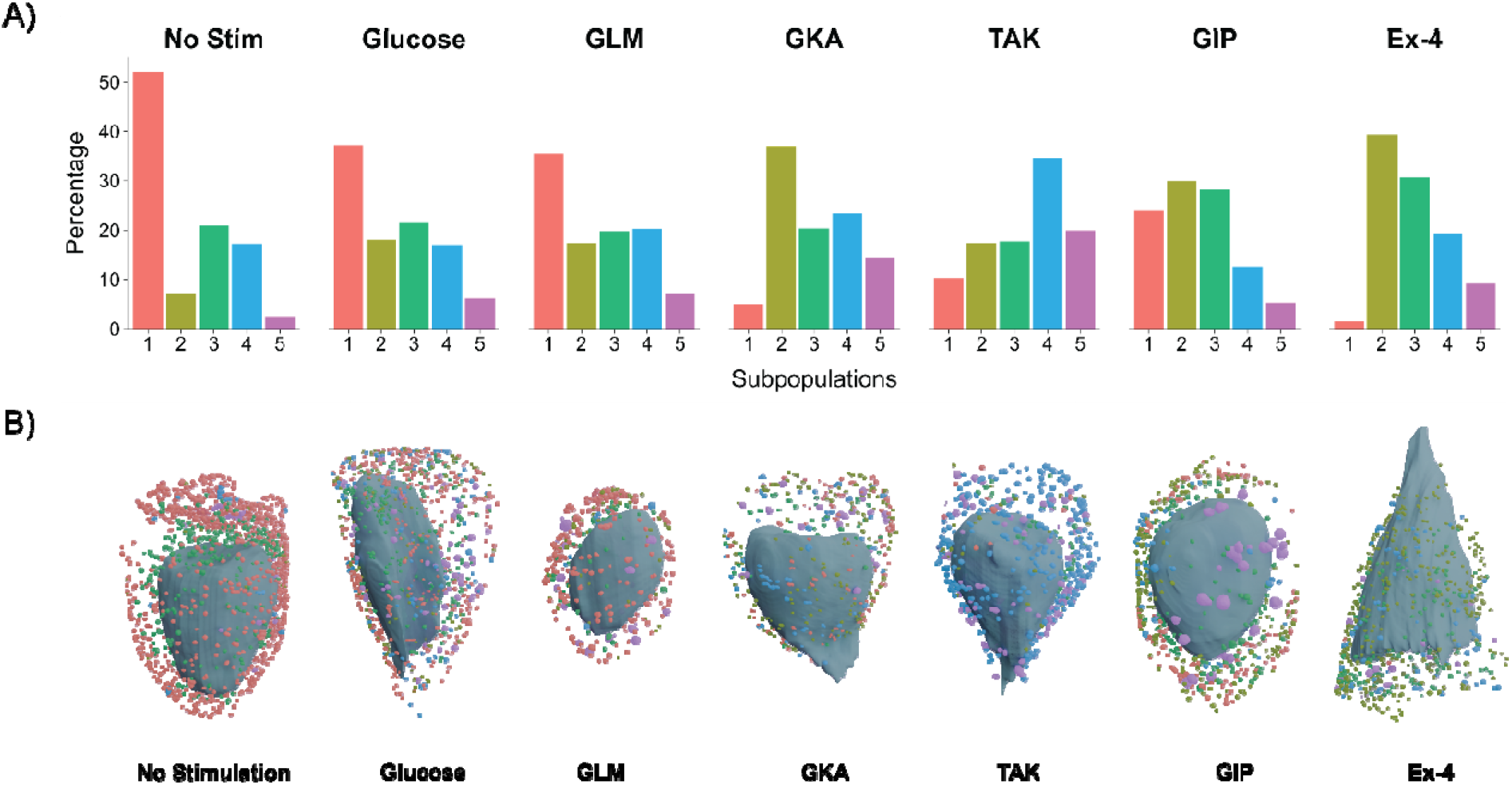
ISG Subpopulation shifts associated with insulin secretory stimuli. (A) Percentage of ISG subpopulations within each stimuli condition. Each condition subpanel shows percentages normalized to the total number of ISGs in that condition, summing to 100%. (B) Blender renderings of representative cells in each stimulation condition. Every cell contains a mask of the nucleus, ISGs, and PM. ISGs are colored according to subpopulation identity. GLM: Glimepiride, GKA50: glucokinase activator 50, TAK: TAK-875, GIP: Gastric Inhibitory Polypeptide, Ex-4: Exendin-4.

In the unstimulated condition, the majority of ISGs are immature granules contained in Subpopulation 1. To stimulate INS-1E cells, the glucose condition was used because glucose treatment is the main mechanism for insulin secretion via glucose-stimulated insulin secretion (GSIS). This condition was still majority Subpopulation 1 but also contained a greater proportion of condensed and mature ISGs compared to the unstimulated condition (Figure 3A). Similar ISG subpopulation shifts were also observed using methods that separate organelles based on their electrokinetic properties.^18^ Glimepiride (GLM), in contrast, is an antidiabetic sulfonylurea that increases insulin release through K_ATP_ mediated membrane depolarization.^37^ Due to the mechanism of action of sulfonylureas, we anticipated and observed that they do not increase the proportion of mature ISGs compared to the glucose condition. Insulin secretory stimuli that act on membrane depolarization lead to the secretion of a wide range of ISGs based on their pH profile compared to glucose stimulation.^4^ This could explain the slight differences in subpopulation distribution between these two conditions.

Using glucokinase activator (GKA50), which increases the metabolic activity of glucokinase resulting in increased ATP production,^38^ we observed an increase in the condensed ISG subtype Subpopulation 2 and the highly mature subtype Subpopulation 5. In this condition, the main effect was an increase in dense core formation in both large and small granules. TAK-875 is a G-protein coupled receptor 40 (GPR40) agonist acting on free fatty acid receptor signaling pathways like inositol triphosphate (IP3) and diacylglycerol (DAG) signaling.^39^ TAK stimulation resulted in a large-scale shift to the mature subtypes, Subpopulation 4 and 5. Secretion of Subpopulation 4, a mature and peripheral ISG subtype, would help explain the observed effect of augmented insulin release during the second phase of prolonged insulin secretion upon TAK administration.^40,41^

Incretins are a class of gut hormones responsible for decreasing blood glucose levels by increasing insulin secretion. Here, we use Gastric Inhibitory Polypeptide (GIP) and Exendin-4 (Ex-4, a glucagon-like peptide-1 receptor (GLP-1R) agonist), which both increase insulin secretion via cyclic-adenosine monophosphate (cAMP) signaling.^42^ Despite their similar signaling pathways, the subpopulation profiles of GIP and Ex-4 are different. The main distinction is between the relative ratios of Subpopulations 1 and 2. Subpopulation 1 is nearly absent and Subpopulation 2 is notably enriched in the Ex-4 condition, while Subpopulation 1 is still a major component in the GIP treated cells. The intracellular insulin to proinsulin ratio is higher in Ex-4 treated INS-1E cells relative to GIP treated cells.^24^ Therefore, Ex-4 could significantly enhance the ISG maturation conversion process in a distinct manner from other drug stimuli.

### Heterogeneity of Docked ISGs

The readily releasable pool of ISGs is often interpreted to be the subset of ISGs primed and docked for release at the plasma membrane (Figure 4A,B).^2,43,44^ In this context, “docked” granules are attached to the plasma membrane via the SNARE complex.^45^ We sought to understand if there was meaningful heterogeneity in this docked pool of ISGs beyond its exocytotic definition by employing a clustering strategy.^46^ To combine the mixture of continuous information such as LAC mean and categorical information such as docked or undocked, we used the Gower’s distance metric (Figure S8A).^47,48^

**Figure 4.**
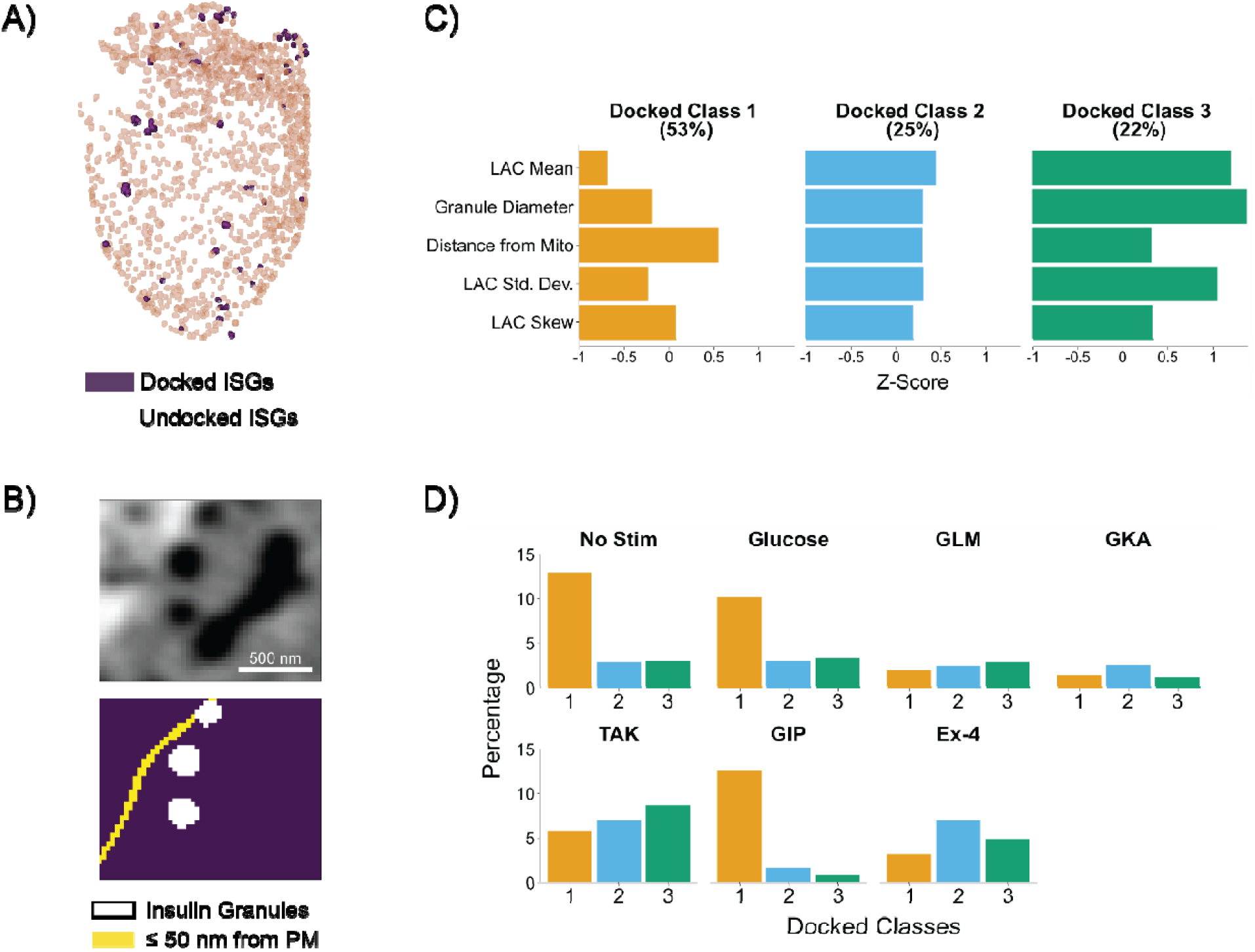
Classifying variants of docked granules. (A) INS-1E cell rendering, with docked and undocked ISGs shown in different colors. Docked ISGs were about 6% of all total ISGs in our dataset. (B) SXT projection image closeup and the same area with the ≤ 50 nm distance from PM region colorized. The ISGs are colored white, while the docked region is colored yellow. Top center ISG is docked at the PM. (C) Profile plot of docked ISG classes. Most docked granules have low LAC. Clusters are organized numerically from lowest to highest LAC. (D) Docked classes organized by condition. These percentages correspond to percent of total overall docked ISGs, since the number of docked ISGs per condition is important to visualize in this analysis.

After performing k-medians clustering on the ∼36,000 ISG dataset using Gower’s distance, we found three classes of docked ISGs (Figure 4C, Figure S8B).^49^ We find a straightforward interpretation of the docked classes, with an increase in ISG LAC mean, LAC standard deviation, and granule diameter from Classes 1 to 3. Over 50% of all docked ISGs were in Docked Class 1, reflecting the fact that most docked ISGs originate from conditions where the INS-1E cells are treated with stimuli that do not increase maturity. Similar to Subpopulation 5 from the previous analysis (Figure 2B), Docked Class 3 is distinguished by elevated LAC mean, diameter, and LAC standard deviation.

When docked ISG pools are categorized according to experimental condition, distinct profiles emerge (Figure 4D). The composition of the unstimulated and glucose condition profiles were similar to each other, with the majority of both being composed of Docked Class 1. This aligns with our expectation for the unstimulated condition, where many ISGs would be primed for release at the PM. Due to the membrane depolarization mechanism of GLM, very few docked ISGs were found. The GKA treated cells have few docked ISGs, suggesting that this stimuli mechanism may alter the dynamics of secretion and replenishment of newly docked ISGs at a 30 minute time point relative to glucose stimulation alone. The GIP condition was primarily composed of Docked Class 1, potentially reflecting an increase in new ISGs that are trafficked to the cellular periphery. In contrast, the TAK and Ex-4 stimulation conditions result in an increase in Docked Classes 2 and 3. While ISGs from TAK stimulated cells are relatively large, increasing the amount of potential surface area that can be in contact with the PM, ISGs from the Exendin-4 condition with generally smaller diameters still display increased PM contacts. Therefore, these findings are not solely due to increased ISG size and point to more dense, mature ISGs ready for release in the second phase of insulin secretion.

### ISG Subpopulations in Primary β-Cells

To evaluate the ISG subpopulation clustering process in a more biologically relevant model, ISGs from mouse primary β-cells were used. When we repeated the analysis on six unstimulated and two Ex-4 stimulated β-cells, totaling to ∼18,000 ISGs, we found 6 clusters using clustering stability analysis (Figure S9A). The distribution and values of unnormalized parameters was similar between the INS-1E and primary cells to allow direct comparison of subpopulations (Figure S10). ISG Subpopulations 1, 2, and 3 in INS-1E cells (Figure 2B) appear to correspond to Primary Subpopulations 1, 4, and 3 in β-cells based on similarities between the z-scores of the five primary ISG parameters (Figure S9A). In contrast, Primary Subpopulation 5 ISGs have large diameters and high LAC standard deviations similar to INS-1E Subpopulation 5 but have comparatively lower LAC means. These variations in biophysical ISG classification may reflect the differences in GSIS response between primary β-cells and insulinoma INS-1E cells.^50^

When primary ISG subpopulations are segregated by condition (Figure S9B), we observe a subpopulation shift that validates results from the INS-1E cells (Figure 3A). Following Ex-4 stimulation, there is a reduction in the proportion of lower LAC mean ISG subpopulations. Additionally, there is an increase in Primary Subpopulation 4, which are smaller, condensed ISGs. These observations can be visually demonstrated with cellular renderings of an unstimulated and Ex-4 stimulated primary cell (Figure S9C). Overall, this data displays similar trends to the INS-1E cells but suggests there may be slight variations in biochemical content between the two cell types.

### Volumetric EM facilitates a complementary perspective of ISG subpopulations

To further test the generalizability of this analysis pipeline, we also analyzed a volumetric EM dataset. While this method lacks the quantitative biochemical density information of SXT, more subcellular structures can be visualized in volumetric EM due to the resolution and membrane contrast principle of this method (Figure 5A). An important metric captured by EM is the distance of ISGs from the Golgi apparatus, which plays a central role in ISG biogenesis and subsequent maturation.^51^ Additionally, completely immature ISGs not easily detected by SXT may be visualized with EM. In Müller et al., the authors used 3D focused ion beam scanning electron microscopy (FIB-SEM) to map seven mouse primary β cells (3 low glucose, 4 high glucose for 1 hour) and described a variety of subcellular organelle rearrangements.^30^ We set out to apply clustering methods on this volumetric EM dataset to identify subpopulations of ISGs that provide a complimentary perspective to the subpopulations found through SXT. In EM, ISGs can be characterized by diameter as well as by their distance from the PM, mitochondria, Golgi, and microtubules.

**Figure 5.**
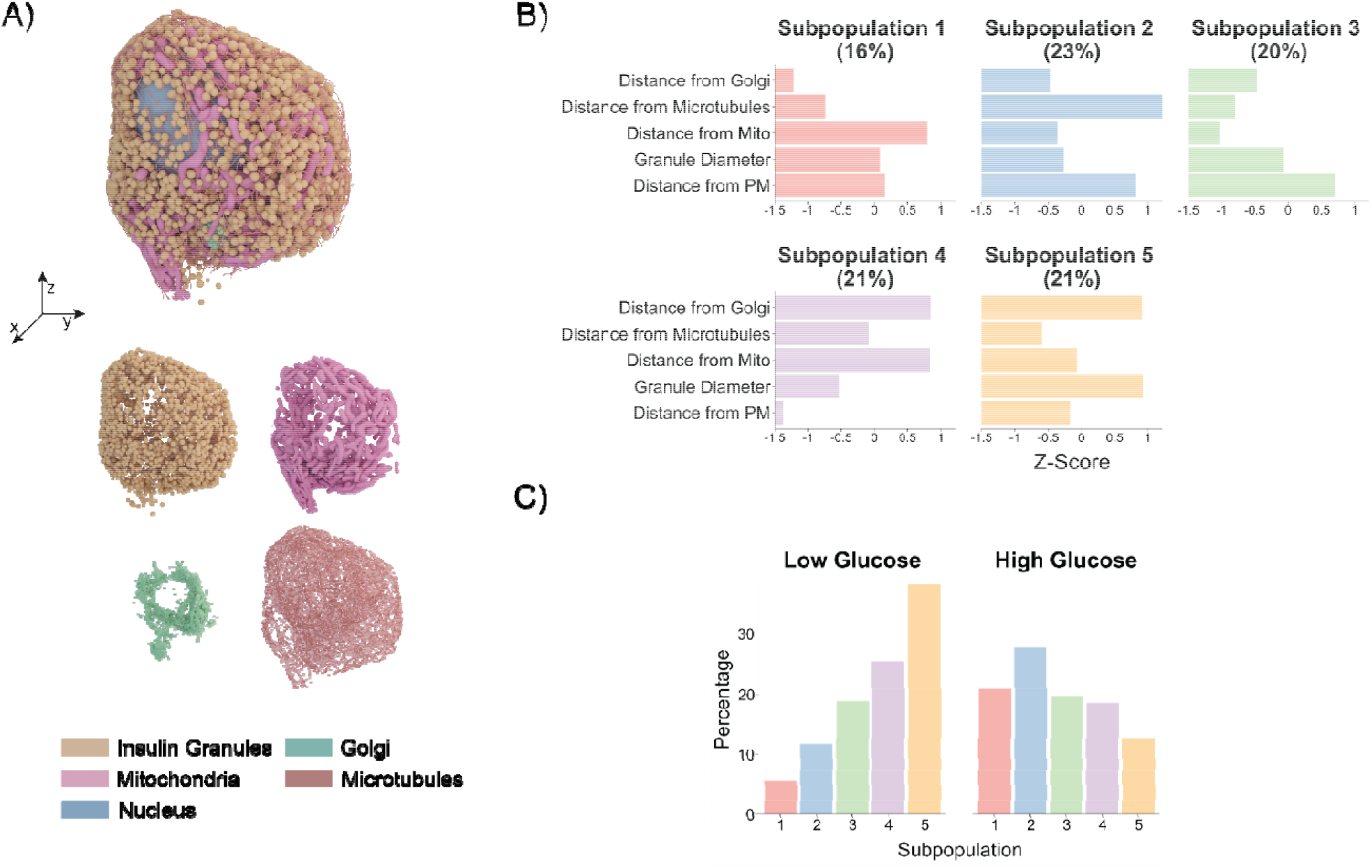
Interpreting Subpopulations in Electron Microscopy data. (A) 3D rendering of a β-cell from the publicly available Müller et al. dataset,^30^ custom-rendered in Blender. Additional organelles like the Golgi apparatus and microtubules can be visualized and quantified using this method. (B) Profile plot of EM ISG Subpopulations. Subpopulations listed in order of distance from Golgi and parameters ranked by importance. (C) EM Subpopulation organized by condition. Total percentage in each subpanel adds up to 100%.

The data parameterization and clustering stability process for continuous parameters was repeated, and 5 stable EM subpopulations were found (Figure 5B). The most important EM-derived ISG parameter for defining subpopulation identity was distance from Golgi (Figure S11A). The correlation matrix of ISG parameters also shows that the features are relatively uncorrelated, although interestingly a slight negative correlation exists between distance from Golgi and distance from plasma membrane (Figure S11B).

Comparison of EM Subpopulations with SXT Subpopulations from primary β-cells (Figure S9A) adds context that neither method could achieve alone. EM Subpopulation 1 are medium sized ISGs near the Golgi apparatus and microtubules, likely representing newly synthesized ISGs. This grouping corresponds to SXT Subpopulation 1, defined by low LAC, linking newly formed ISGs to immature dense core granules. Primary SXT Subpopulation 2 and EM Subpopulation 3 can also be matched together, as both are far from the PM and have average-sized diameters. These characteristics suggest that they are reserve pool ISGs. EM Subpopulation 5 and SXT Subpopulation 5 both consist of large ISGs. Their extended distance from the Golgi and elevated LAC values, respectively, suggest a population of highly mature, dense ISGs.

Additionally, shifts in EM Subpopulations are apparent between two glucose-based conditions (Figure 5C). Subpopulations 1 and 2 increase in the high glucose condition, which indicates newly synthesized ISGs being created at the Golgi and trafficked along microtubules. Interestingly, the high glucose condition has a lower proportion of Subpopulation 5 compared to low glucose. This reduction may reflect the depletion of mature ISGs through insulin secretion under prolonged stimulation. Although this trend differs from our SXT findings, it may be explained by the difference in treatment duration, as these cells were exposed to glucose for 1 hour compared to 30 minutes in the SXT experiments.

## DISCUSSION

Here, we used a variety of clustering methods to delineate subpopulations of ISGs. We find that different secretory stimuli change the proportions of ISG subpopulations in INS-1E cells and in primary β-cells. Applying different clustering approaches such as Gower’s distance provided new perspectives into ISG heterogeneity. Integrating EM and SXT information created complimentary ISG subpopulation definitions, demonstrating the utility of clustering methodology on whole cell data.

The advent of whole cell datasets at mesoscale resolution makes it possible to inventory the vast community of organelles within a single cell.^17^ While advancements in SXT and vEM technologies have expanded our ability to map cells in their entirety, analytical strategies capable of fully exploiting this data remain underdeveloped. To bridge the disconnect between experimental sophistication and downstream analysis, we adopted the conceptual framework of morphomics - the systematic, unbiased, and quantitative exploration of cellular structure.^52,53^ Inspired by other “omics” methodologies, we use clustering approaches to identify subpopulations of ISG, enabling us to explore patterns of cellular organization in a way that parallels developments in spatial proteomics,^54^ connectomics,^34^ and computational representations of cells.^55^ We envision that our framework will be a broadly applicable strategy for organelle subpopulation discovery and the analysis of cellular systems, generalizable across whole cell methods and cell types.

In this work, we wanted to systematically expand our ability to identify populations of ISGs beyond the canonical readily releasable pool and reserve pool. Likely, our observations here overlap with other ISG subpopulation identities based on molecular composition and pH. A disease-orientated study found decreased granule diameter and increased amounts of sphingomyelin and cholesterol from ISGs labeled with synaptotagmin-7 (syt7), a calcium binding protein that promotes fusion of granules to the PM.^12^ In SXT, these characteristics would correspond to a population characterized by decreased diameter but appreciable LAC content, which we observe in Subpopulation 2 and 3 (Figure 2B). Another modality characterizing ISG subpopulations by pH and spatial location in the cell revealed that ISGs far from the PM exhibit increased acidity.^4^ This pool of ISGs might correlate to Subpopulation 5, which is characterized by high LAC and a localization also further away from the PM. The increased acidity in these ISGs would result in a more processed insulin maturation state, leading to the formation of a dense insulin core.

This broad analytical framework also provides a basis for developing new therapies aimed at enhancing the quality of ISGs secreted by β-cells by enriching for specific ISG subpopulations.^15^ Traditionally, therapeutic development for diabetes has focused primarily on increasing overall insulin secretion; however, certain pharmacological interventions that chronically upregulate insulin release may lead to β-cell exhaustion and long-term treatment failure.^56^ Clustering methodologies, used in diverse fields to detect outliers or target particular consumer segments,^49^ offer an analogous approach to determine if therapeutics may affect a given ISG subpopulation of interest. For instance, the shift to smaller but biochemically dense ISG subpopulations during Ex-4 treatment (Figure 3A, Figure S9B) would help explain the beneficial effects of GLP-1R agonists without necessarily contributing to β-cell exhaustion and depletion.

When using SXT to image INS-1E and primary β-cells, inherent constraints in spatial resolution and contrast likely limits the detection of ISGs, especially immature populations.^31^ For instance, β-cells imaged using vEM tend to have many more ISGs per cell compared to β-cells visualized with SXT.^30^ Therefore, when unsupervised clustering is performed on ISGs from SXT-imaged cells, the subpopulations may be biased towards mature ISG states. While ISGs obtained from vEM data may be more representative of the immature ISG population, the distinction between immature and mature granules is less clear without the LAC parameter. To improve the description of ISG maturity in EM-based datasets for subpopulation analysis, a categorical classifier based on ISG core shapes could be used. While whole cell, mesoscale datasets provide a plethora of information per cell, the overall number of cells is limited in this study. In total 44 INS-1E (SXT), 8 primary β-cells (SXT), and 7 primary β-cells (vEM) for a total of 59 whole cells were analyzed. While sufficient to describe several ISG states, imaging and processing more cells per condition could validate our findings and potentially enable us to discover new ISG subpopulations with novel biological roles. Advances in vEM image acquisition speed and automated segmentation using machine learning could increase throughput in large-volume EM, while increased accessibility of SXT technology could increase the sample size of cells to study.^59,60^ Several docked ISG states were described in our study (Figure 4). While this analysis is useful for distinguishing between docked ISG characteristics, we cannot rule out if the ISGs in this region are ready for release, involved in kiss and run secretion, or tethered to the membrane.^2^ Future work using cryo-ET to distinguish between these different states may provide additional insights into docked ISG heterogeneity.

Future applications of this method could explore the role of disease progression on ISG subpopulations. Disease state β-cells are known to be in an altered metabolic state and have impaired insulin processing,^13,57^ which would likely impact the distribution of ISG subpopulations. Characterizing shifts in ISG subpopulations or new subpopulations that arise as the result of disease states would highlight mesoscale level changes to organelle ensembles caused by diabetes. Correlative investigations using spatial proteomics would integrate cellular context with established ISG subpopulation biomarkers. ISG subpopulations have been described using a wide variety of protein markers related to age or release potential,^2,12^ yet their localization in the cell is not well established. Such investigations may provide a “Rosetta stone” for correlating different ISG subpopulations together. Additionally, utilizing a high-resolution technique like Cryo-ET would provide insights into the molecular milleu of ISGs. Observations on subpopulations with higher ISG LAC and diameter could be paired with detailed information about the protein and lipid makeup of individual ISGs.^58^

## RESOURCE AVAILABILITY

### Lead contact

Requests for further information and resources should be directed to and will be fulfilled by the lead contact, Kate L. White.

### Materials availability

This study did not generate new reagents.

### Data and code availability

- All reconstructed tomograms are available upon request from the corresponding author.
- All original code has been deposited at GitHub at https://github.com/kvn42999/Classifying-Biophysical-Subpopulations-of-ISGS and is publicly available as of the date of publication.
- Any additional information required to reanalyze the data reported in this paper is available from the lead contact upon request.

## ACKNOWLEDGMENTS

We thank Carolyn Larabell and Mark Le Gros at the National Center for X-ray Tomography (NCXT) for use of the XM-2 SXT microscope and cryogenic confocal fluorescent microscope at the Advanced Light Source (LBNL, Berkeley). Research at NCXT is supported by the NIH (NIGMS P30GM138441 to Carolyn Larabell) and the DOE Biology and Environmental Research Project (DE-AC02-05CH11231 to Carolyn Larabell). Thanks to Axel Ekman for his support with tomogram reconstruction. We would like to thank Jitin Singla for his insightful discussions about performing ISG subpopulation analysis. Thanks to Janielle Cuala and Tvisha Singh for helping with islet dissociation and Jeffrey Velasquez for his molecular biology assistance. Thanks to Peter Arvan for providing the hPro-CpepSfGFP mouse line. Funding from the National Institute of General Medical Sciences of the National Institutes of Health under award number R35GM154893 and the USC Bridge Institute at USC helped financially support this work.

## AUTHOR CONTRIBUTIONS

Conceptualization, K.C. and K.L.W.; methodology, K.C, A.D., V.L., and K.L.W.; Investigation, K.C., A.D., V.L., and R.V.; writing—original draft, K.C. and K.L.W.; writing—review & editing, K.C, A.D., R.V., V.L., and K.L.W.; funding acquisition, K.L.W.; resources, K.L.W.; supervision, V.L. and K.L.W.

## Methods Cell Culture

Full sample preparation details can be found in Deshmukh et al., 2025.^24^ Briefly, a rat insulinoma β-cell line, INS-1E,^61^ was used for experiments. Cells were cultured in T75 flasks with Optimized RPMI media supplemented with 10% FBS, 1× Pen/Strep, and 0.05 mM β-mercaptoethanol and seeded at a density of 4.0×10^5^ cells/cm^2^. All cells were grown at 37°C, in a 5% CO_2_ atmosphere.

## Animal Model

All animal studies involving SXT were conducted using procedures approved and conducted per Institutional Animal Care and Use Committee (IACUC) guidelines at the University of Southern California (Animal Use Protocol #21120). Primary β-cells were collected from the C57BL6/J mouse model expressing human proinsulin tagged with C-peptide-Superfolder Green Fluorescent Protein (CpepSfGFP), provided by Peter Arvan (University of Michigan).^62^ Mice were 2-3 months of age for experiments.

## Cell Stimulation

For stimulation experiments, the cells were starved for 30 minutes in a low glucose (1.1 mM) buffer and then co-stimulated with 25 mM glucose and a specific stimulation condition for 30 minutes (Glimepiride: 100 nM, Glucokinase activator 50: 100 nM, TAK-875: 10 µM, Gastric Inhibitory Peptide: 10 nM, and Exendin-4: 10 nM). The no stimulation condition was in the starvation treatment only. After stimulation, cells were centrifuged, resuspended in 1x PBS, and then loaded into custom made glass capillaries with tip diameters ranging from 6-10 µm. The capillaries were then rapidly plunged into liquid nitrogen-cooled liquid propane and stored in liquid nitrogen.

## Primary Cell Preparation

Islets were extracted from mice pancreases, dissociated into single cells, and then transported to the SXT microscope facility. To distinguish β-cells from other islet cell types, a cryogenic confocal fluorescence microscope was used to correlate the β-cell GFP signal with the X-ray absorption signal in the glass capillaries.^21^

## SXT Data Collection

SXT imaging was performed at the National Center for X-ray Tomography located at Lawrence Berkeley National Laboratory (LBNL). Projection images were collected at 517 eV using the SXT microscope XM-2, which was equipped with a 50 nm resolution objective lens.^72^ To minimize radiation damage, cell samples in capillaries were exposed to a steady stream of liquid nitrogen-cooled helium gas. The projection images were sequentially collected around a 180° axis of rotation (capillary axis) in 2° increments, with an exposure time of 350 ms for each projection. Drifts in the capillary image were corrected automatically by the data acquisition software and 3D image reconstruction was achieved by the software package AREC3D.^65^ LAC values were calculated by normalizing the intensity value of each pixel by the pixel area, which allows for comparison of LAC values across different cells.^22^

## Tomogram Segmentation

Tomograms were segmented using the ThermoFisher Amira 2021.2-2023.1 software. ISGs were identified based on their shape, size, and characteristic LAC value range.^19–21^ While lipid droplets may be another circular, dense organelle, their large size (> 600 nm) and LAC (mean LAC >0.55 µm-1) allows us to differentiate them from ISGs. Mitochondria and the nucleus were segmented based on characteristic morphology and LAC. The initial segmentation for the plasma membrane (PM) was obtained through the ACSeg 3D UNET model on Biomedisa,^64^ followed by manual editing on Amira. In particular, the “magic wand” tool on Amira was used extensively to segment ISGs and mitochondria in a semi-automatic manner.

## Data Parameterization and Preprocessing

ISG parameters were obtained using a custom Python (version 3.12.3) code that used the raw LAC file and organelle masks of cells as inputs (Figure S1A). Python packages cv2 and cc3d were used to process organelle masks. Based on literature values for ISG, ISG with diameters greater than 600 nm were excluded from the dataset.^73^ Distance metrics were calculated via the Euclidean distance transform (EDT) and were normalized to the maximum membrane EDT to account for variations in cell shape and size.

To compare each parameter on the same relative scale for clustering, every ISG parameter was standardized using z-score normalization. LAC Skew and Kurtosis were calculated to further describe the distribution of LAC values in ISGs (Figure S1B). To account for the highly right skewed distribution of distance measurements (Figure S1C), an additional log normalization was applied beforehand to ISG distances from the PM and mitochondria.^49^ The final list of parameters used to describe ISGs is LAC mean, LAC standard deviation, LAC skew, LAC kurtosis, diameter, and distances from the PM and mitochondria. While other ISG parameters such as LAC max and granule volume were calculated, only one representative feature out of several correlated features was used in the final analysis to accurately depict trends in ISG subpopulations (Figure S2).

## ISG Clustering Analysis

Clustering Analysis was performed using a custom R (version 4.4.1) code (Figure S1A). Clustering stability analysis was performed using the flexclust R package with the bootFlexclust function.^32,67^

A stable cluster configuration was considered to be the maximum possible number of clusters that had an average Jaccard index over 0.75 (relative to bootstrapped clustering results).^74–76^ Once a stable number of clusters was found, the clustering assignments were calculated using the kcca k-medians function also found in the flexclust package. To calculate feature importances for ISG parameters, each parameter was individually permuted across all ISGs, and clustering was repeated on the resulting permuted dataset. Then, the Rand index was calculated between the original and permuted cluster partitions. The lower the Rand index, the more important a feature. Uniform manifold approximation and projection (UMAP) dimensional reduction was performed using the R umap package.^69^

For analyses combining a mixture of continuous and categorical data, Gower’s distance metric was calculated using the gower.dist function in the StatMatch R package.^71^ To categorize ISGs in contact with the PM, a distance of less than or equal to 50 nm between the ISG and PM was used.^77^ When calculating Gower distances, the weight of the binarized PM distance metric was varied. The final PM distance weight was chosen to yield a clustering configuration suitable for characterizing docked ISG heterogeneity. Visualization of clustering results was performed using ggplot2.^70^

## 3D FIB-SEM Analysis

Visualization of clustering results Exact details on the EM imaging methods and conditions used on β-cells can be found in Müller et al., 2020.^30^ Data preprocessing and clustering methods are the same as in the main methods section, except with the omission of LAC data and inclusion of distance from Golgi and from microtubules.

## Blender Cellular Renderings

Binary masks of cell organelles and components, particularly ISGs, mitochondria, the nucleus, and the plasma membrane, are converted from TIFF to STL using Python libraries Trimesh and skimage, and imported into Blender for smoothening and material rendering. The ISG binary mask is analyzed using connected-components-3d (cc3d) to correlate cluster assignments and location imported from derived datasets. This information is used to generate voxel-to-vertex visuals in 3D space through Blender scripting in Python and geometry nodes. Vertices are individually colored to match the entire ISG while maintaining organic, native structure.

