## Supplementary Figures for "Classifying Biophysical Subpopulations of Insulin Secretory Granules using Quantitative Whole Cell Structure Analysis"

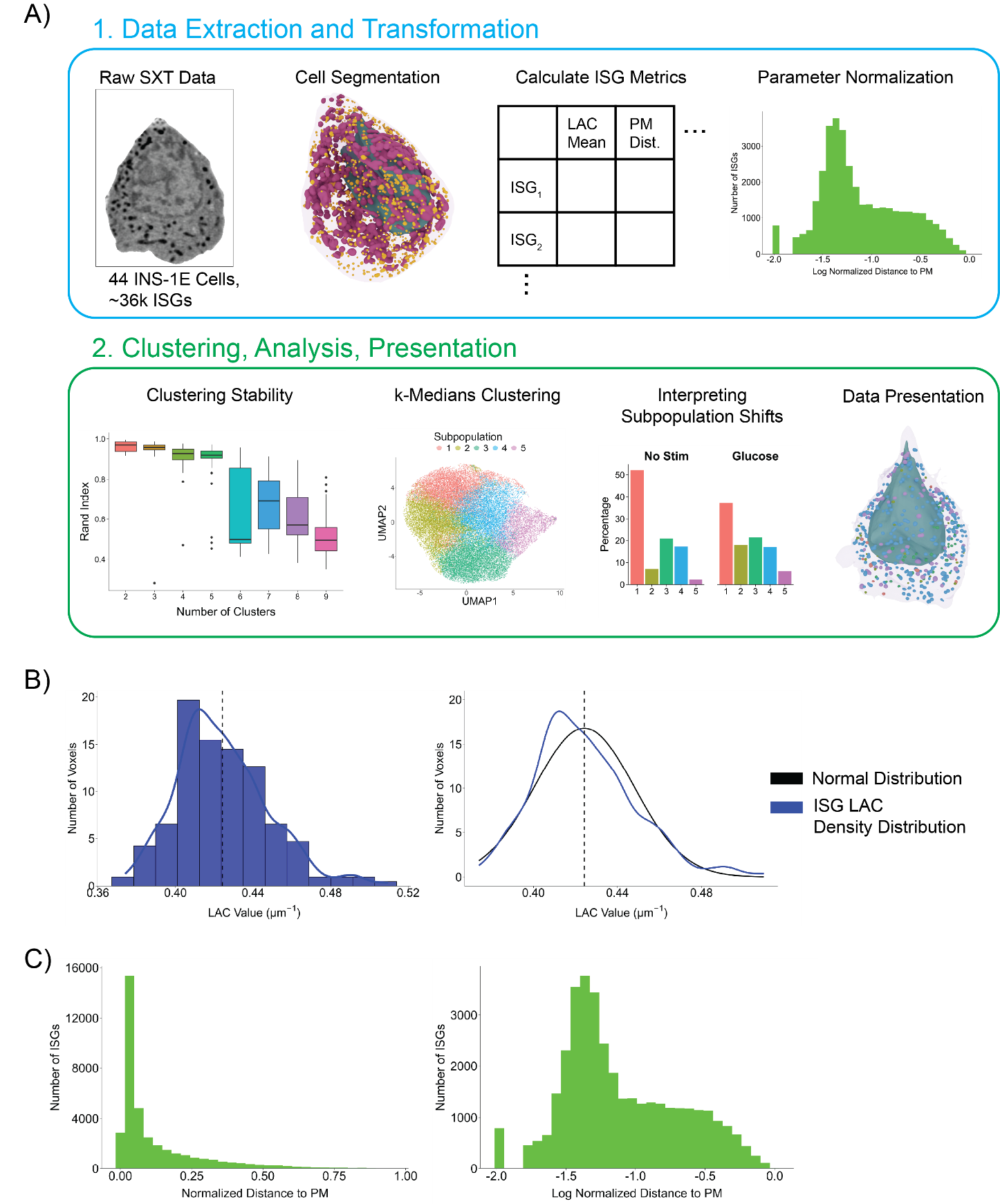

**Figure S1:** Workflow diagram and calculation of ISG parameters. (A) Workflow of ISG clustering analysis pipeline. Raw INS-1E SXT data was segmented using Amira into various organelle masks. Using the LAC cellular image data and organelle masks, individual ISG metrics were calculated. ISG metrics underwent various data normalization steps before use in clustering. Clustering stability analysis using our normalized ISG metrics guided the k-medians clustering process. Once final ISG subpopulation assignments were created, subpopulation shifts under different experimental conditions could be determined. Using Blender, INS-1E cell renderings were created with ISGs colored by their subpopulation identity. (B) Histogram of the LAC values in a single ISG shown in blue, with the mean LAC value of the ISG displayed as a dashed line and distribution density estimate shown by the blue line. LAC density estimate is compared to a normal distribution relative to the mean LAC value. The discrepancy between the two distributions allows for the calculation of parameters such as LAC Skew and Kurtosis. (C) Histograms showing the ISGs distances from the PM (maximum PM EDT normalized) in green. Log normalization mitigates the highly skewed distribution of the distance measurements, allowing for more meaningful comparisons to other ISG parameters through z-score normalization.

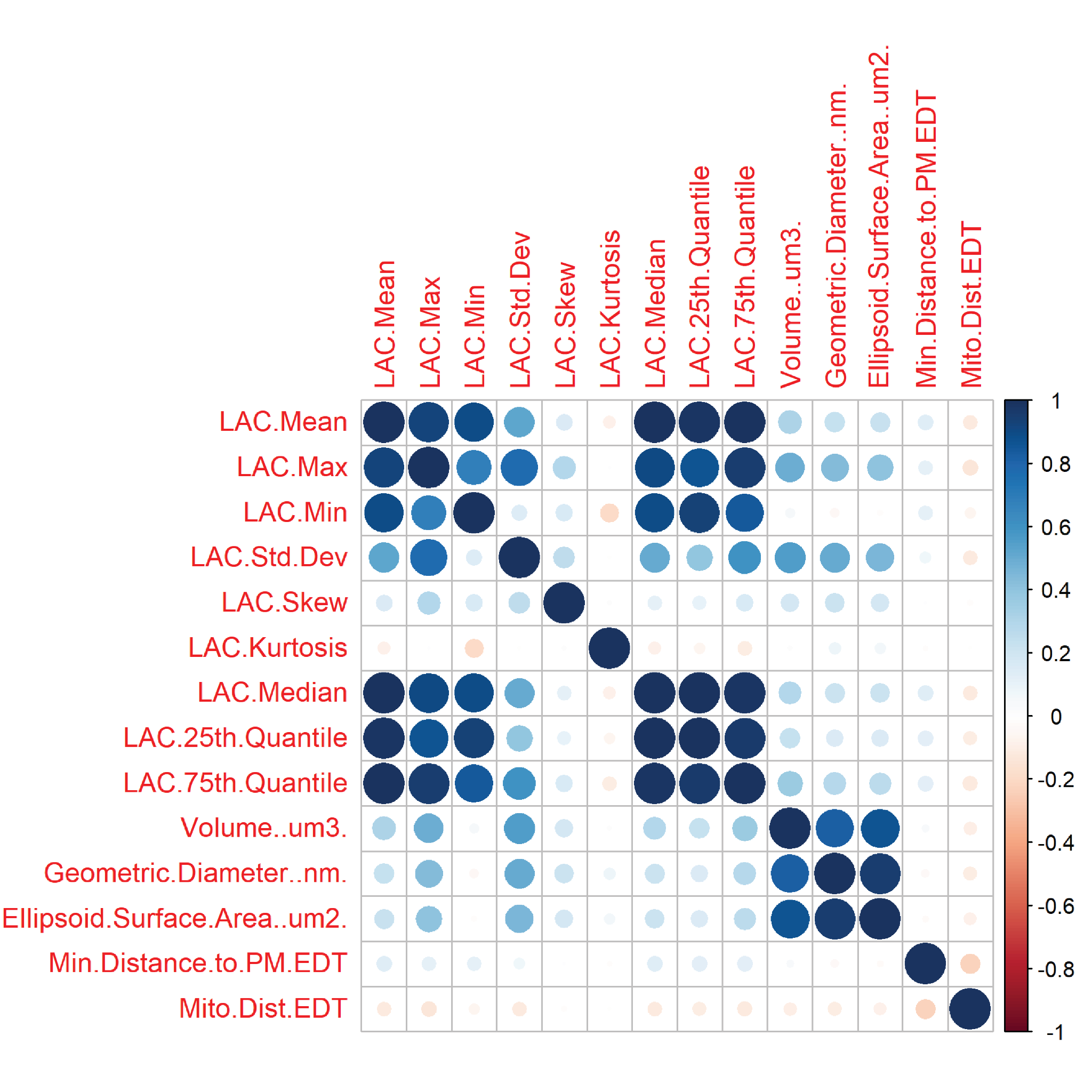

**Figure S2:** Correlation plot of ISG Parameters. Values of the Pearson correlation coefficient are shown in the legend on the right, where negative correlation is shown in red and positive correlation is shown in blue. The degree of correlation is also indicated in the figure by the area of the circle. Most parameters have weak correlation with each other, with the exception of ISG size parameters and measures of the central tendency of LAC distributions. Therefore, ISG Diameter and LAC Mean were used to represent these trends respectively.

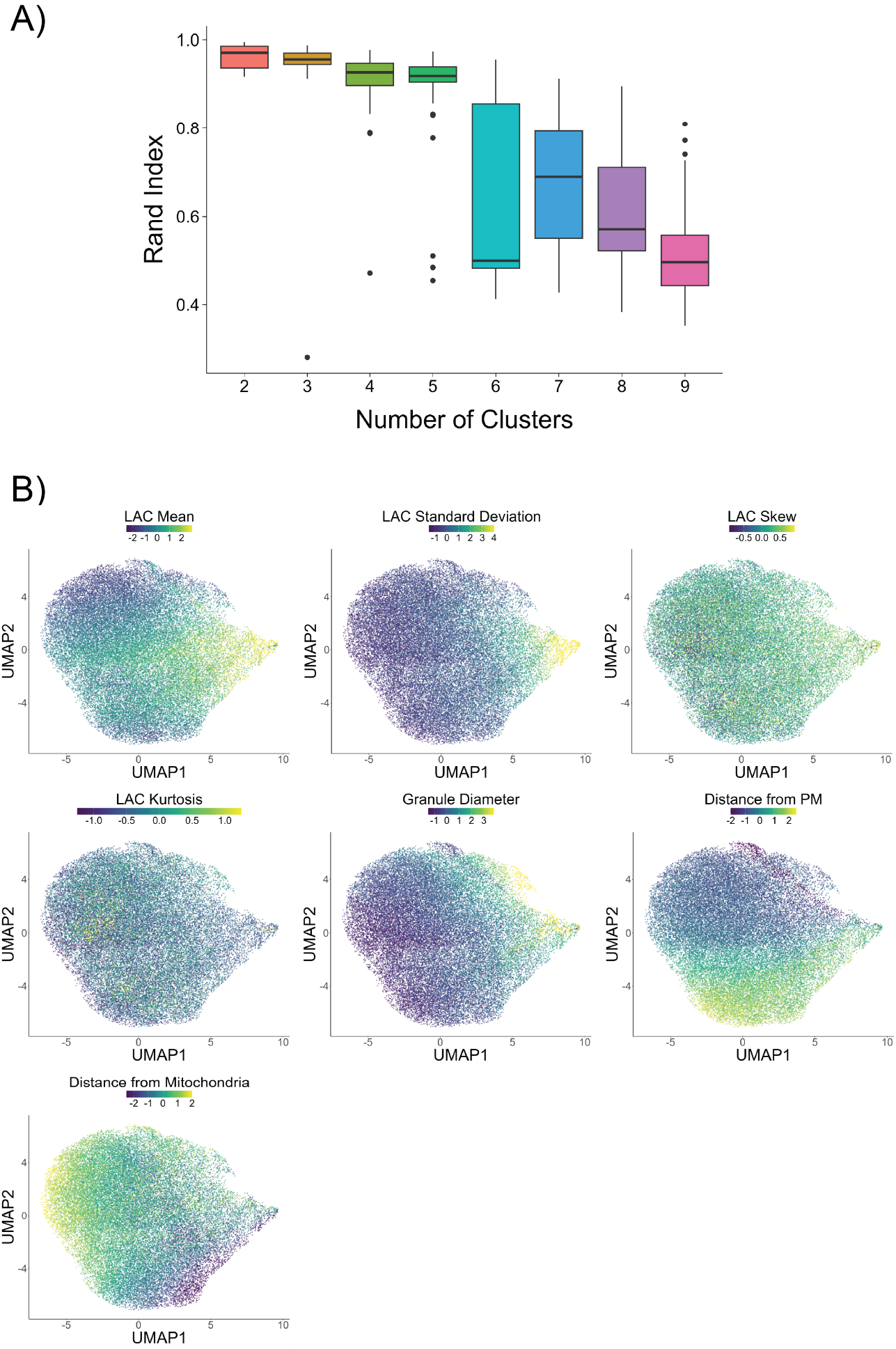

**Figure S3:** Assessment of ISG Subpopulations. (A) Figure displaying the number of stable clusters using clustering stability analysis. To assess clustering stability, 100 bootstrapped datasets were generated, and the Rand index was calculated to measure the consistency of clustering results across varying numbers of clusters (*k*) in each bootstrapped dataset. Five clusters were chosen for downstream analysis because this configuration captures the greatest amount of robust, explainable heterogeneity. (B) UMAP diagrams of ISGs colored by LAC Mean, LAC Standard deviation, LAC Skew, LAC Kurtosis, Granule Diameter, Distance from PM, and Distance from Mitochondria.

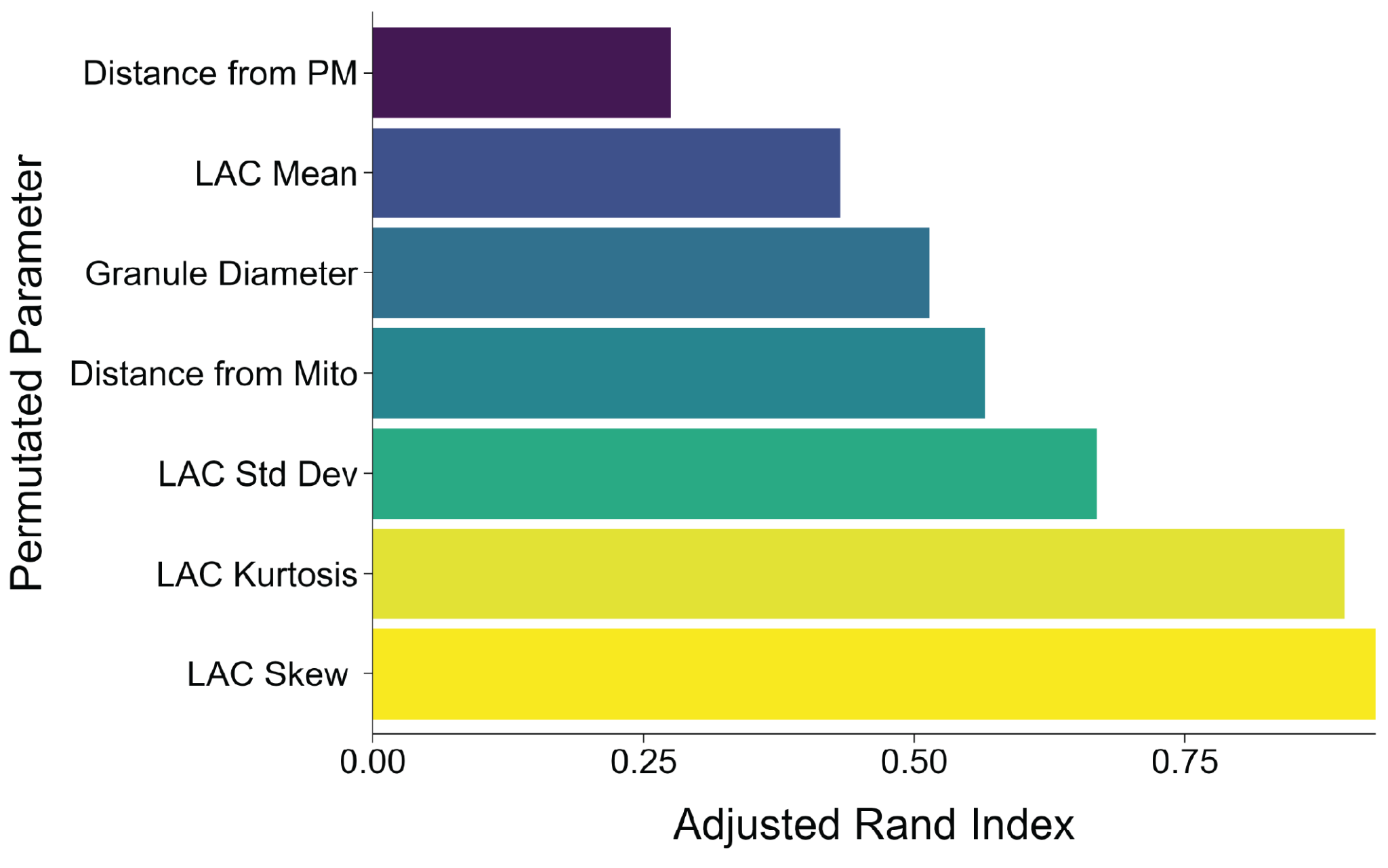

**Figure S4:** Determination of the most important ISG parameters influencing subpopulation identity. To determine importance, the values of a specific ISG parameter were permuted across all ISGs. Clustering was performed on this permuted dataset and then compared to the original cluster assignments using the Rand index. This process is repeated 100 times, and the maximum Rand index was used as the final measure of subpopulation identity importance.

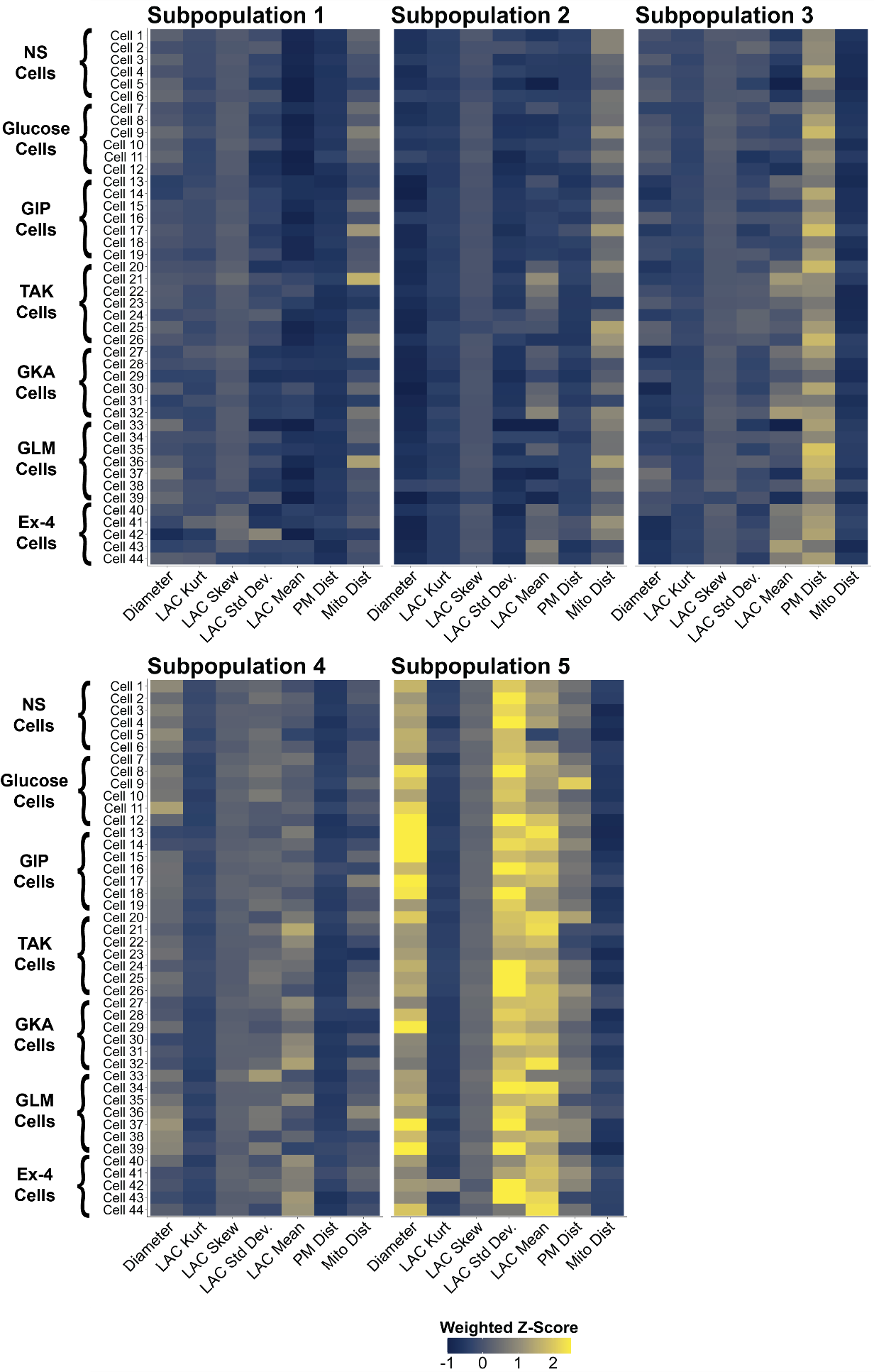

**Figure S5:** Visual assessment of how effectively each ISG subpopulation, formed from ISGs aggregated across multiple cells, reflects trends present in individual cells. Rows list INS-1E cells (grouped by experimental condition), while columns list ISG parameters. Each heatmap box is colored according to the average weighted z-score of a given parameter across all the ISGs within a single cell. If a subpopulation represents individual cells convincingly, each column should have similar trends.

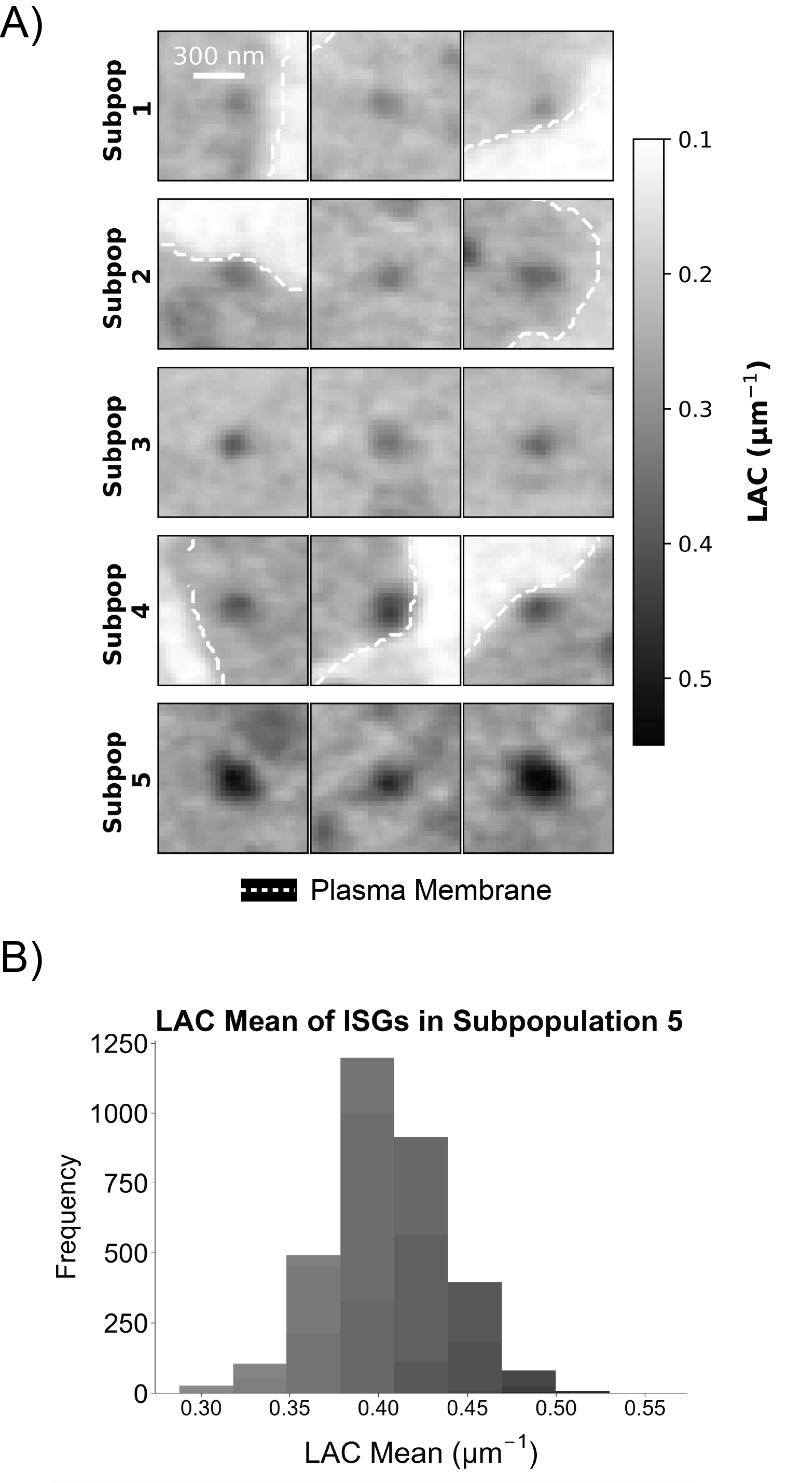

**Figure S6:** Biochemically dense material in ISG Subpopulation 5. (A) ISG gallery with the maximum displayed LAC increased to 0.55 μm^-1^, in contrast to the 0.40 μm^-1^ LAC maximum used in Figure 2C. Dense cores are especially present in ISG Subpopulation 5. (B) Histogram of ISG LAC means in Subpopulation 5.

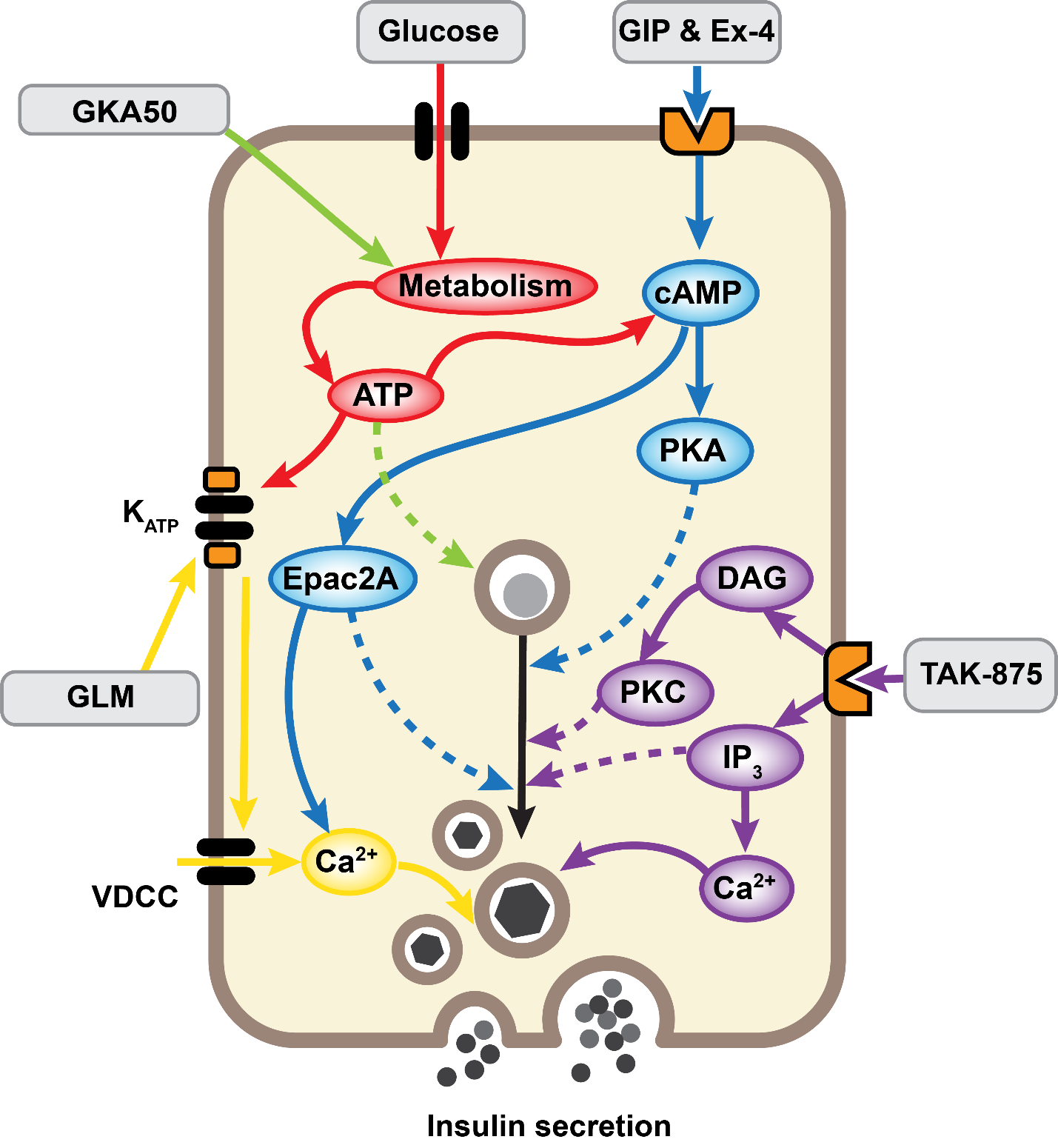

**Figure S7:** Different cellular signaling pathways activated by insulin secretory stimuli. Figure modified from Deshmukh et al, 2025. Metabolic signaling shown in red, sulfonylurea signaling in yellow, incretin signaling in blue, and GPR40 agonist signaling in purple. These pathways are believed to have differing effects on ISG maturation, as represented by the conversion of an ISG containing diffuse proinsulin into an ISG with a dense, insulin core. GLM: Glimepiride, GKA50: glucokinase activator 50, GIP: Gastric Inhibitory Polypeptide, Ex-4: Exendin-4.

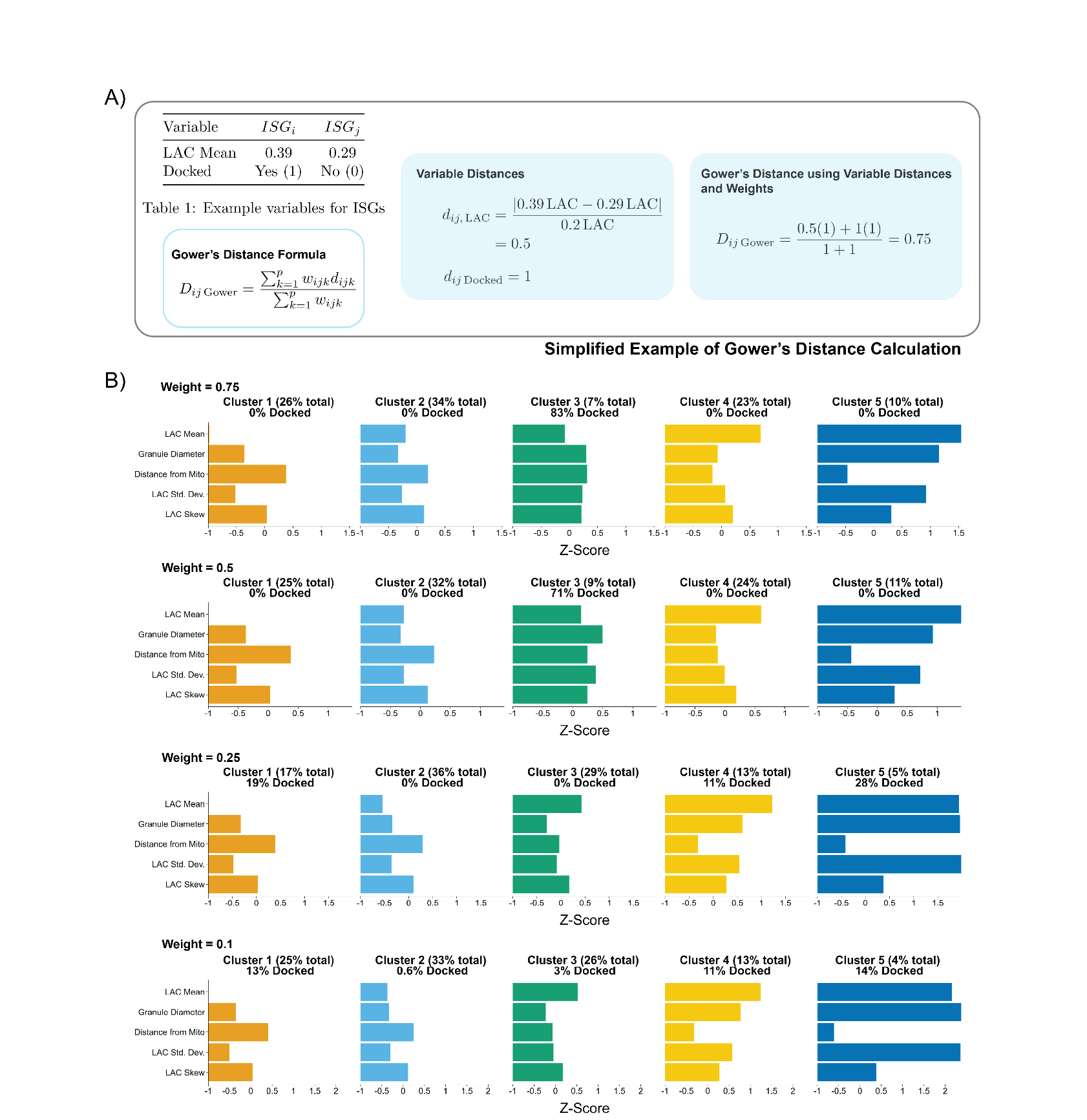

**Figure S8:** Gower’s distance calculations and evaluation. (A) Brief sample calculation of Gower’s distance using two example ISGs. The table shows two ISGs with LAC mean and docked features. Below is the generalizable Gower’s distance equation. Variable distances are calculated for each variable; for continuous variables such as LAC mean range normalization is used, while matching coefficients are used for binary variables. Based on variables weights *w* (all *w* = 1 in this example), Gower’s distance can be calculated between two ISGs. (B) Gower distance clustering results using different feature weights for the binarized ISG docked/not docked feature. To determine an appropriate binary PM distance weight for Gower’s distance, we tested multiple weight values. Clustering at weight = 0.75 or 0.5 places all docked ISGs into one subpopulation. Clustering at weight = 0.25 results in a relatively stable clustering configuration, while clustering at weight = 0.1 does not show a significant difference from clustering at weight = 0.25. Therefore, clustering using weight = 0.25 was used for the final downstream analysis since it was appropriate for understanding docked ISG heterogeneity.

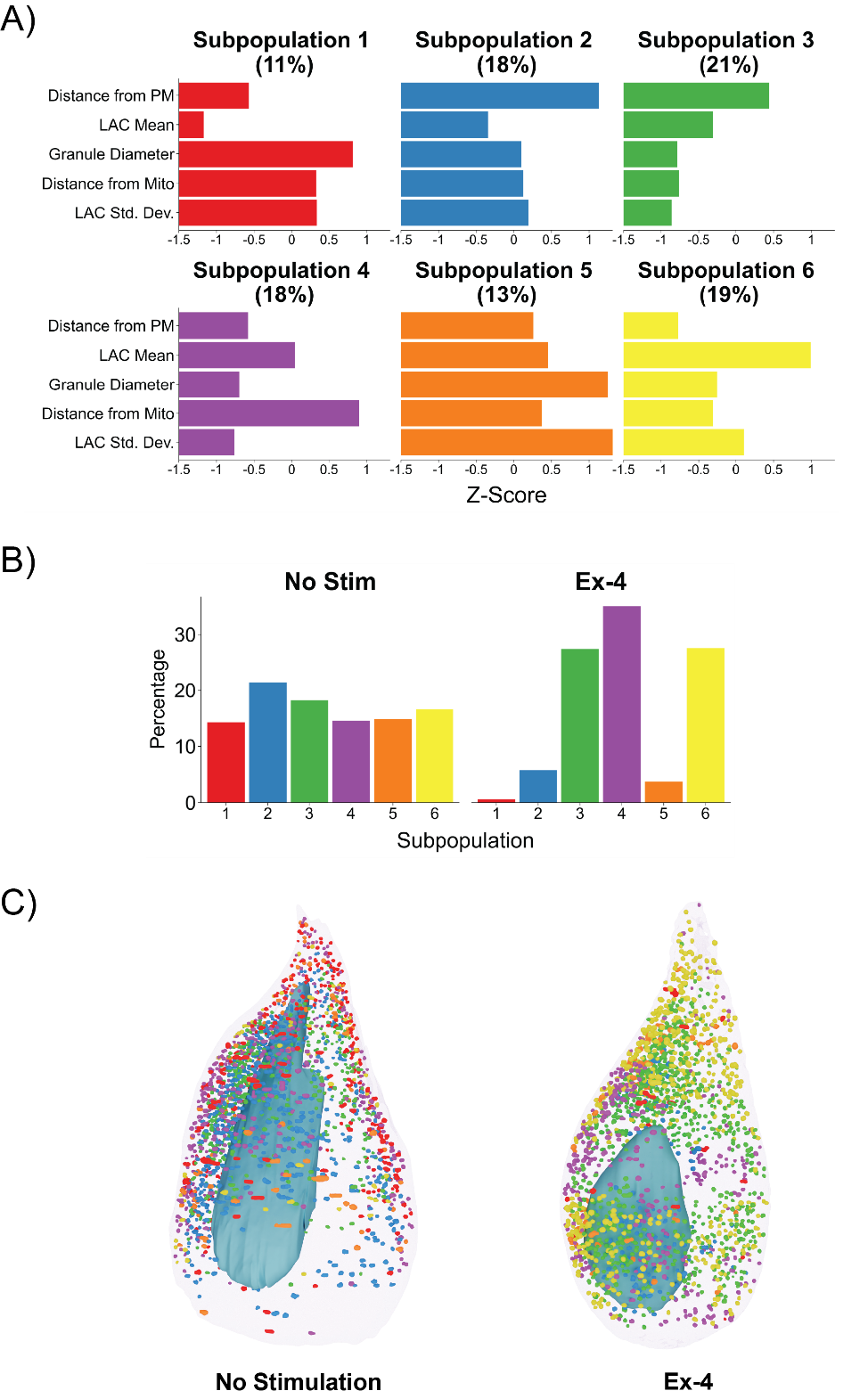

**Figure S9:** Subpopulations in primary mice β-cells. (A) ISG subpopulation profile plot obtained through cluster stability analysis of the primary β-cell dataset. (B) Percentage of ISGs within the unstimulated and Ex-4 conditions normalized by the number of ISGs per condition. Subpopulations drastically change between the two conditions. (C) Primary β-cell renderings made in Blender. The unstimulated cell is on the left while the Exendin-4 cell is on the right. ISGs are colored by cluster identity.

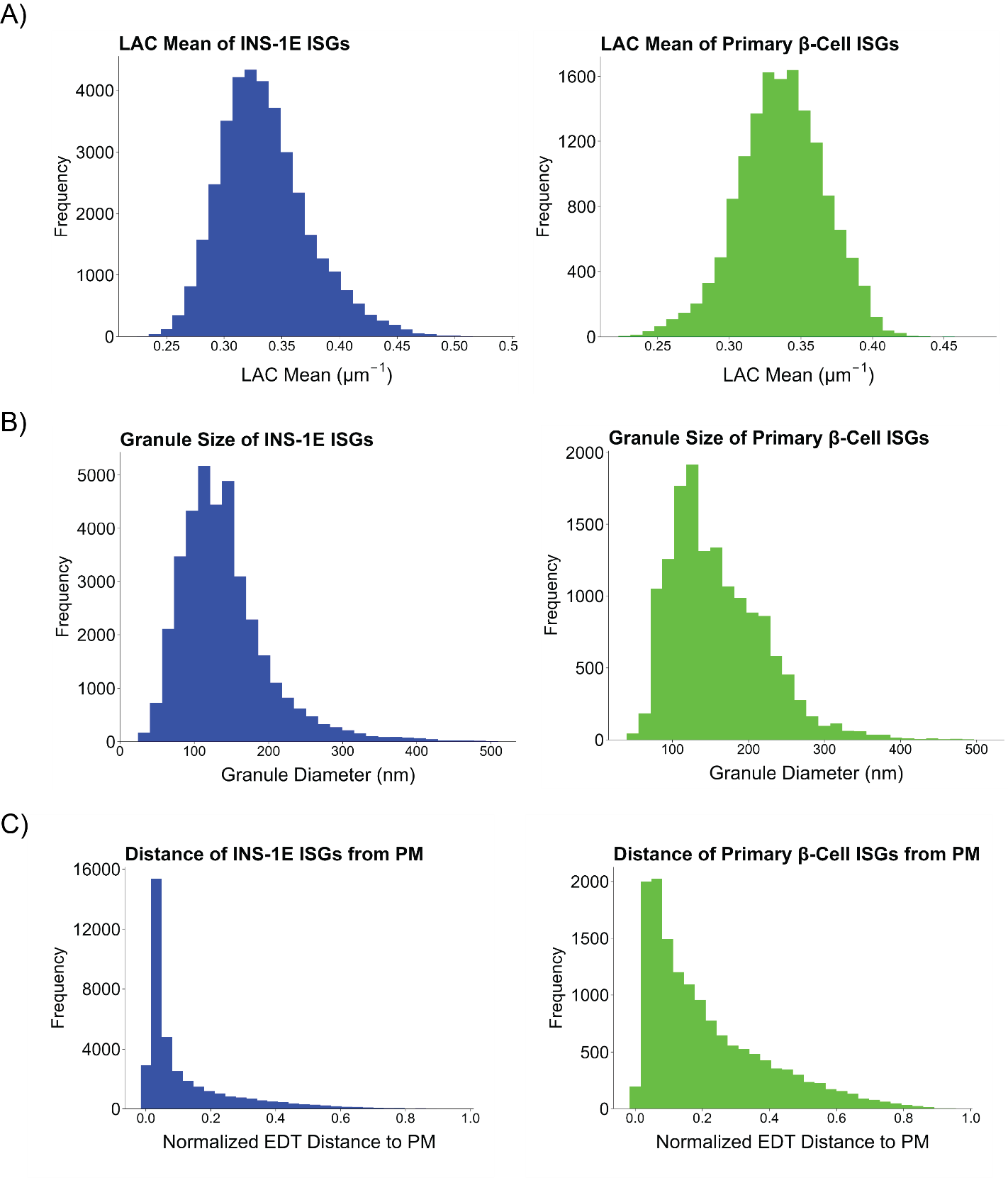

**Figure S10:** Comparison of ISG features in INS-1E and primary β-cells. (A) Histogram displaying ISG LAC Means in INS-1E (blue) and primary β-cells (green). (B) Histogram displaying ISG Diameters in INS-1E (blue) and primary β-cells (green). (C) Histogram displaying ISG distance from the PM in INS-1E (blue) and primary β-cells (green).

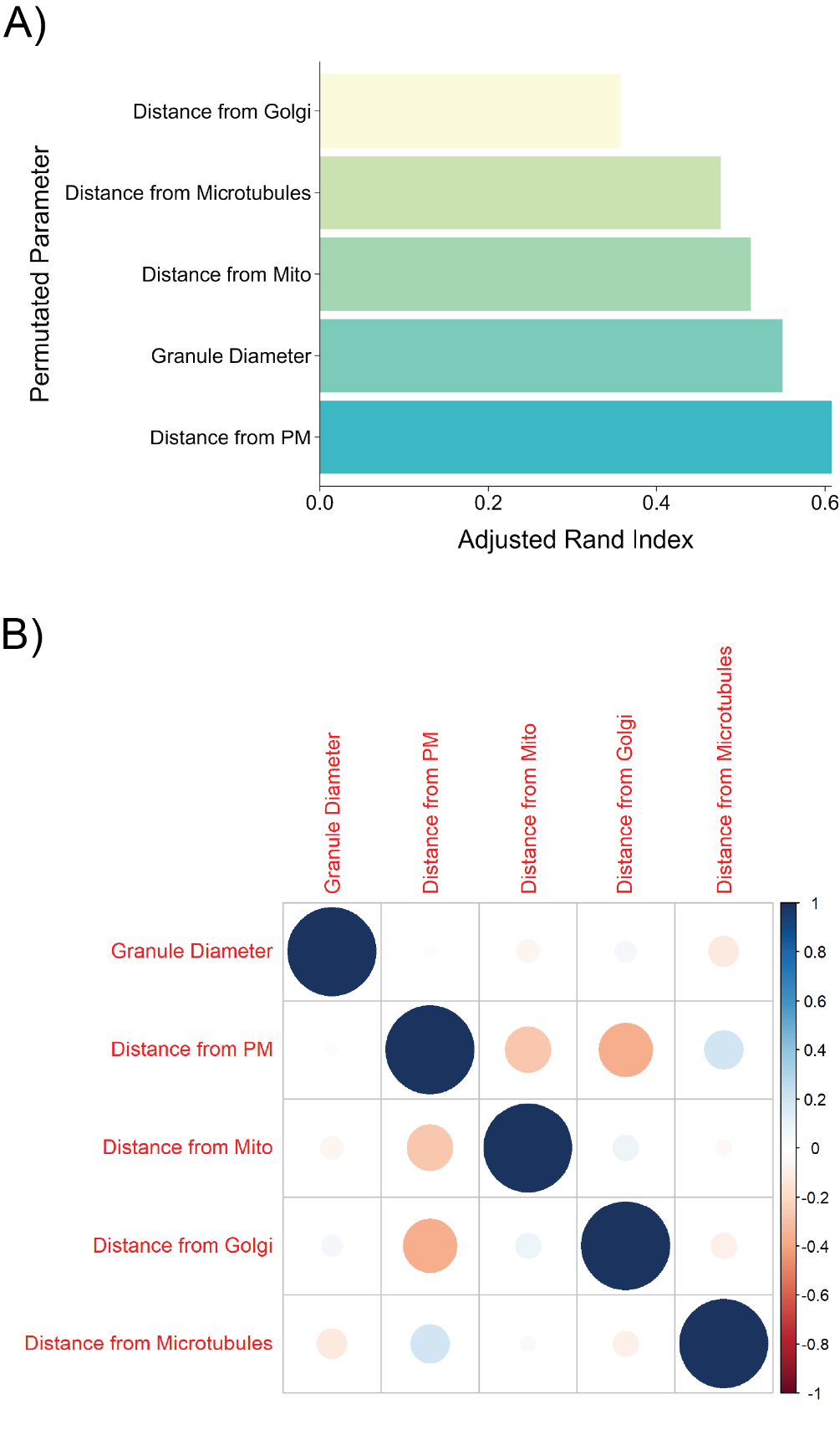

**Figure S11:** Evaluation of ISG features in the FIB-SEM data of primary mice β-cells. (A) Feature importance plot of ISG parameters influencing subpopulation identity. Brief overview of importance in this context explained in Figure S4. (B) Correlation plot of ISG Parameters. Pearson correlation coefficient indicated by color and area of circles.
